# Successional Stages in Infant Gut Microbiota Maturation

**DOI:** 10.1101/2021.06.25.450009

**Authors:** Leen Beller, Ward Deboutte, Gwen Falony, Sara Vieira-Silva, Raul Yhossef Tito, Mireia Valles-Colomer, Leen Rymenans, Daan Jansen, Lore Van Espen, Maria Ioanna Papadaki, Chenyan Shi, Claude Kwe Yinda, Mark Zeller, Karoline Faust, Marc Van Ranst, Jeroen Raes, Jelle Matthijnssens

## Abstract

**Background:** Disturbances in the primary colonization of the infant gut can result in life-long consequences and have been associated with a range of host conditions. Although early life factors have been shown to affect the infant gut microbiota development, our current understanding of the human gut colonization in early life remains limited.

To gain more insights in the unique dynamics of this rapidly evolving ecosystem, we investigated the microbiota over the first year of life in eight densely sampled infants (total number of samples, n=303). To evaluate gut microbiota maturation transition towards an adult configuration, we compared the microbiome composition of the infants to the Flemish Gut Flora Project population (n=1,106).

**Results:** We observed the infant gut microbiota to mature through three distinct, conserved stages of ecosystem development. Across these successional gut microbiota maturation stages, genus predominance was observed to shift from *Escherichia* over *Bifidobacterium* to *Bacteroides*. Both disease and antibiotic treatment were observed to be associated occasionally with gut microbiota maturation stage regression, a transient setback in microbiota maturation dynamics. Although the studied microbiota trajectories evolved to more adult-like constellations, microbiome community typing against the background of the Flemish Gut Flora Project (FGFP) cohort clustered all infant samples within the (in adults) potentially dysbiotic Bact2 enterotype.

**Conclusion:** We confirmed similarities between infant gut microbial colonization and adult dysbiosis. A profound knowledge about the primary gut colonization process in infants might provide crucial insights into how the secondary colonization of a dysbiotic adult gut can be redirected.

## BACKGROUND

Development of a stable adult large-intestinal microbiota sets off with the primary colonization of the infant gut. Disturbances in initial colonization or ecosystem maturation can potentially result in life-long consequences and have been associated with a broad range of host conditions, including inflammatory bowel disease[1], asthma[2], and type I diabetes[3]. Although early life factors such as birth mode and diet have been shown to affect the development of the infant gut microbiota[4, 5], our current understanding of the human gut colonization in early life still remains limited. Microbiome monitoring efforts combining high sampling frequency with prolonged longitudinal design would enable gaining more insights in the unique dynamics of this rapidly evolving ecosystem. Here, we investigated microbiome variation over the first year of life in eight densely sampled infants, analysing on average 38 samples per participant (total number of samples, n=303). We observed the infant gut microbiota to mature through three distinct, conserved stages of ecosystem development. Across these successional gut microbiota maturation stages, genus predominance was observed to shift from *Escherichia* over *Bifidobacterium* to *Bacteroides*. A stable, reproducible order of successive colonization could be established at genus level across the BaBel infants. Both disease and antibiotic treatment were observed to be associated occasionally with gut microbiota maturation stage regression – a transient setback in microbiota maturation dynamics. Although the studied microbiota trajectories evolved both in terms of richness and composition to more adult-like constellations, microbiome community typing against the background of the n=1,106 Flemish Gut Flora Project population cohort clustered all infant samples within the (in adults) potentially dysbiotic Bact2 enterotype. While these observations reflect incomplete microbiota maturation within the first year of life, the suggested parallel between primary succession as observed in the healthy infant’s gut and secondary colonization upon ecosystem disruption could inform novel bio-therapeutic approaches based on sequential recolonization of a dysbiotic community.

## RESTULTS AND DISCUSSION

An increasing number of diseases is being linked to gut dysbiosis. This state - characterised by a low diversity and high abundance of facultative anaerobic bacteria in adults – also resembles the microbiome composition of healthy infants[6]. A profound knowledge about the primary gut colonization process in infants, going from (nearly) sterile at birth towards a diverse and healthy gut microbiota later in life, might provide crucial insights into how the secondary colonization of a dysbiotic adult gut can be redirected.

### The colonization process in the healthy infant gut happens through distinct stages of ecosystem development

Setting out to map gut microbiota maturation dynamics in eight vaginally delivered, healthy infants from Belgium (BaBel cohort), we analysed faecal microbiome profiles of a core dataset of 142 samples collected on predefined time points distributed over the first year of life (from the 159 samples at predefined time points, 17 were excluded based on reported disease signs; Supplementary Table 1a; Supplementary Figure 1), complemented with 144 post-hoc selected samples associated with clinically relevant events such as disease/drug treatment. Applying Dirichlet Multinomial Mixtures (DMM) modelling on the microbiome profiles, we screened for sub-communities among the infants’ microbiomes. Grouping samples potentially originating from a same community through probabilistic modelling, DMM-based stratification of microbiome data reproducibly identifies community constellations across datasets without making any claims regarding the putative discrete nature of the strata detected[7]. In the present dataset, community typing revealed the presence of four compositionally distinct clusters or gut microbiota maturation stages, with only one of them restricted to a single individual (Supplementary Table 1b; Figure 1a; Supplementary Figure 2). Three out of four maturation stages (labelled A, B, and C) comprised almost exclusively samples originating from seven out of eight individuals, reflecting conserved, structured microbiome maturation rather than inter-individual variation. Although time-of-transition varied between individuals (Figure 1b), maturation stage A-C succession revealed a strong temporal organization following a conserved pattern across infants (n=7, Kendall test, Kendall’s w corrected=1, p-value=5e-4; Figure 1a), aligning with an overall increase in microbiome richness (comparison maturation stage A with B and B with C, n=[182:176], Kruskal-Walis [KW] with post-hoc Dunn test [phD], r=[-0.35:-0.60], FDR<0.05; Supplementary Table 1c; Figure 1c). Highest richness values were however noted for the divergent D maturation state (comparison maturation stage A, B and C vs D, n=[127:119:121], KW with phD test, r=[-1:-0.72:-0.18], FDR<0.05; Supplementary Table 1c; Figure 1c) – only observed in infant S011 and not linked to temporal variation. Focusing on differences in microbiota composition between the gut microbiota maturation stages, we found maturation stage A to be dominated by *Escherichia* spp. (Figure 1d). Compared to both B and C, maturation stage A was characterized by higher proportional abundances of not only *Escherichia*, but also *Staphylococcus*, *Enterococcus*, *Enterobacter*, and *Lactobacillus*, among others (n=303, KW with phD test, r>0.3, FDR<0.05; Supplementary Table 1d; Supplementary Figure 3). The reported top five (in terms of effect size) of maturation stage A-associated genera consist exclusively of facultative anaerobic genera, reflecting the higher oxygen levels present in the infant gut shortly after birth[8]. Maturation stage B was dominated by bifidobacteria (Figure 1d), with *Bifidobacterium* being the only genus that was proportionally more abundant in B when compared to both A and C (n=303, KW with phD tests, r>0.4, FDR<0.05; Supplementary Table 1d; Supplementary Figure 3). At the end of their first year of life, all studied infants eventually reached the *Bacteroides*-dominated C maturation stage (Figure 1d). With respect to both A and B, the higher richness of the C maturation stage was reflected in higher proportions of a broad range of bacteria, including butyrate-producing taxa[9] such as *Anaerostipes*, *Faecalibacterium* and *Roseburia* (n=303, KW with phD, r>0.3, FDR<0.05; Supplementary Table 1d; Supplementary Figure 3).

**Figure 1.**
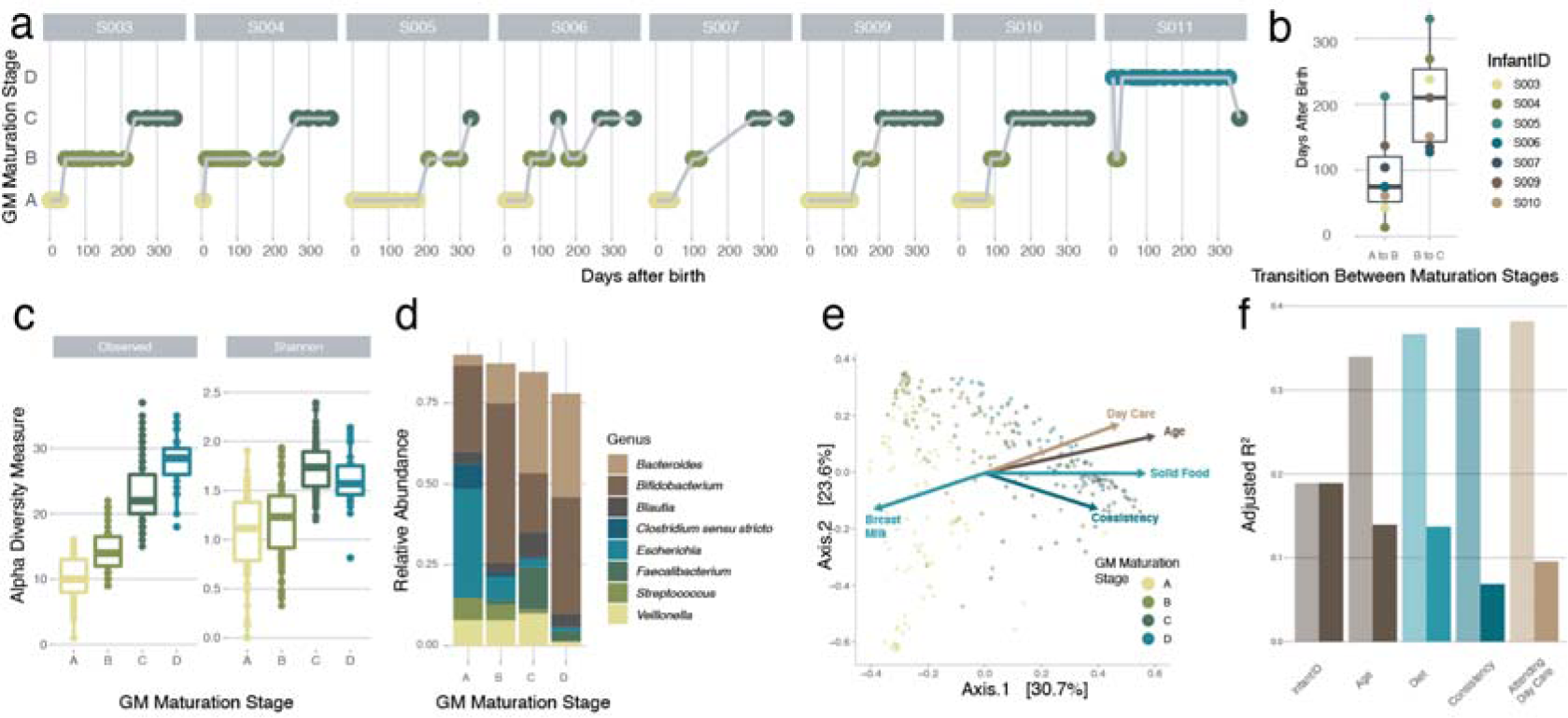
Detailed overview of the colonization process in the healthy infant gut at genus level. (a) Overview of the gut microbiota maturation stage succession of the samples of all the infants over time, coloured by the assigned gut microbiota maturation stages determined using the DMM approach (calculated on all samples (n = 303) and shown here for the samples at predefined time points where the infants were not sick (n = 142)). (b) Variation in timing of transition between the gut microbiota maturation stages in the different infants. The body of the box plots represent the first and third quartiles of the distribution and the median line. (c) Alpha diversity measures (observed ASV richness and Shannon diversity) of the samples within every gut microbiota maturation stage, increasing from A-C (comparison gut microbiota maturation stage A with B and B with C, n = [182:176], post-hoc Dunn test [phD], r = [- 0.35:-0.60], FDR < 0.05.) (d) Mean relative abundance of the most common genera at every gut microbiota maturation stage. (e) Principle coordinate analysis (PCoA, Bray- Curtis dissimilarity) representing genus-level microbiome variation in our infant cohort (n = 299). Dots represent one sample and are coloured by their assigned gut microbiota maturation stage. The arrows represent the effect size and direction of the post-hoc fit of variables significantly associated to microbiota compositional variation (univariate dbRDA, infant ID was excluded for clarity). (f) Covariates with non-redundant explanatory power on the genus level ordination, determined by multivariate distance- based redundancy analysis at genus-level (dbRDA, Bray-Curtis dissimilarity, FDR < 0.05). The light bars represent the cumulative explanatory power (stepwise dbRDA R^2^) and the darker bars represent the individual univariate explanatory power of the variables (dbRDA R^2^). Covariates present in less than three infants were excluded.

### Identification of covariates explaining infant gut microbiota variation

To identify covariates of microbiome diversification within the first year of life, we assessed the non-redundant explanatory power of diet, medication, health status, environment, and infants’ specific characteristics, such as having siblings or their blood group, in genus-level compositional variation within the BaBel infants. Beyond inter-individual variation (n=299, multivariate stepwise distance-based redundancy analysis [dbRDA] on Bray-Curtis dissimilarity, R^2^=18.9 %, p.adj=0.002), microbiome composition was significantly associated with age (R^2^=15.0 %), diet (R^2^=2.7 %), stool consistency (R^2^=0.8 %), and attending day care (R^2^=0.8 %; Supplementary Table 1e; Figure 1e,f). Next, we applied a similar approach to assess potential associations between metadata variables and the top 15 most dominant genera (covering in average 92.6 % of samples total abundance) as identified based on their average proportional abundance over all samples (n=299, multivariate stepwise dbRDA with Euclidean distance on composition, constraining for infant ID, FDR<0.05). Beyond inter-individual variation, we found the effect size of diet to exceed the impact of age in 6 out of 15 genera (Supplementary Table 1f; Figure 2a). Among those, we highlight the complex associations between the omnipresent *Bifidobacterium* spp. and changes in infants’ nutrition[3]. While the taxon as a whole was the lowest in the samples where the infant was weaned (Breast-Milk Only:Non-Solid Food (i.e. breast and formula milk or formula only) vs Solid Food, n=[236:185], phD test, r>0.2, FDR<0.05; Supplementary Table 1g), divergent patterns could be observed when zooming in on the two main amplicon sequence variants (ASV) detected (Figure 2c,d): while ASV1 proportions decreased significantly upon weaning (with weaning defined as the first time solid foods are introduced; Breast Milk Only:No Solid Food vs Solid Food, n=[236:185], phD test, r>0.25, FDR<0.05), ASV2 increased substantially after the addition of formula milk to the diet (Breast Milk Only vs No Solid Food:Solid Food, n=[177:236], phD, r>0.3, FDR<0.05; Supplementary Table 1g; Figure 2e,f).

**Figure 2.**
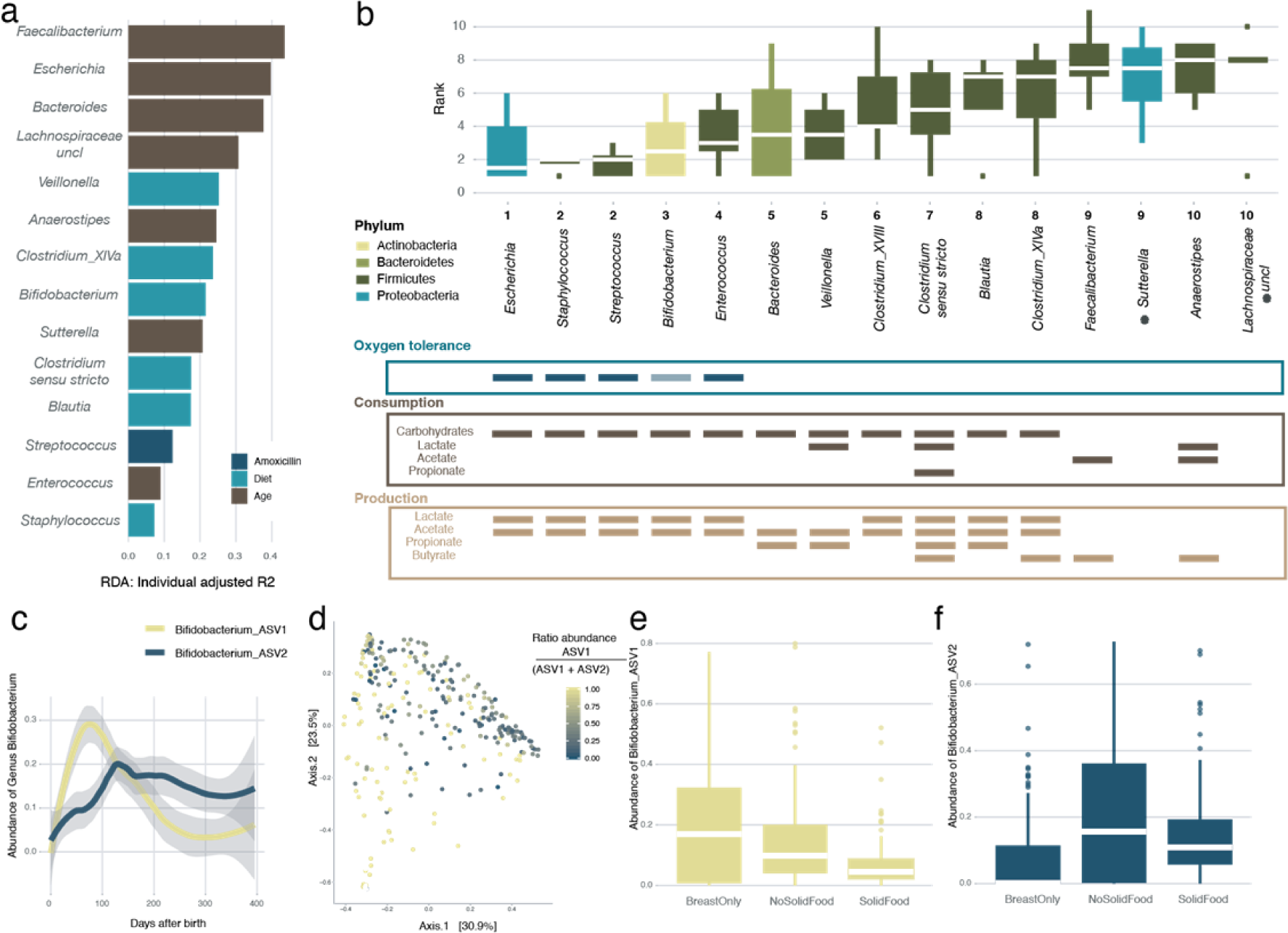
Order of appearance of the most common genera in the infant gut. (a) Overview of the covariates with highest explanatory power for the variation of the top 15 genera in our infant cohort, beyond intra-infant variability (note that for *Clostridium cluster XVIII* no significance was reached). A multivariate redundancy analysis was carried out on the relative abundances of each genus, after constraining for infant ID (multivariate dbRDA, FDR <0.05). The length of the horizontal bars represents the explanatory power of the most significant covariate (stepwise dbRDA R^2^). (b) Order of appearance (presence defined as abundance > 0.5 %) of the top 15 most abundant genera in the infant gut. The boxplots are coloured according to the phylum the genus belongs to. Shown below the boxplots, is the oxygen tolerance of the different genera (note that *Bifidobacterium*, while normally assumed to be strictly anaerobe, is found to be oxygen-tolerant in the human gut[10]), and the consumption and production of different short chain fatty acids (SCFA) by the different genera[11], [12], [9] . The body of the box plots represent the first and third quartiles of the distribution and the median line. The asterisks (*) indicate the genera for which no information was available. (c) The average relative abundances of the different Bifidobacterium Amplicon Sequencing Variants (ASVs) over time averaged over all infants (Loess smoothing). (d) Genus level principle coordinate analysis (n = 299, PCoA, Bray-Curtis dissimilarity), coloured for the ratio of the two most abundant *Bifidobacterium* ASVs. (e) Effect of food on the relative abundance of Bifidobacterium ASV1 showing a higher absence during weaning (Breast Milk Only : No Solid Food vs Solid Food, n = [236 : 185], post-hoc Dunn test [phD] test, r > 0.25, FDR < 0.05). (f) Effect of food on the relative abundance of *Bifidobacterium* ASV2 showing an increase in samples where the infants was having a formula milk-based diet (with or without solid food)(Breast Milk Only vs No Solid Food : Solid Food, n = [177: 236], phD, r > 0.3, FDR < 0.05; Supplementary Table 1g).

### Infant gut microbiota genera appear in a stable, reproducible order

To assess whether microbiota maturation of the infant gut was determined by a series of successional colonization events conserved across individuals, we zoomed in on the genus rather than the community level, investigating the order of appearance of the top 15 most dominant genera within each one-year maturation timeline. Defining appearance as the first occurrence of a genus (relative abundance >0.5 %), we established an appearance ranking for the taxa in each infant. We observed the appearance ranking to be significantly conserved across individuals (n=8, Kendall test, Kendall’s w corrected=0.523, p-value=2.08e-7; Figure 2b). Lowest ranks (*i.e.* primary colonizers) were mainly attributed to genera that have been described as saccharolytic, oxygen-tolerant, and/or lactate- and acetate producing[9–13]. While such taxa can contribute to colonization resistance of the newborns through acidification of the large-intestinal environment[14, 15], they also generate substrates that allow subsequent recruitment of cross-feeders such as *Veillonella* and *Anaerostipes*[16]. Ranks correlated negatively with estimated growth rates, with early colonizers displaying the shortest minimal generation times (n=14, Pearson correlation, r=-0.63, p-value=0.016, Supplementary Figure 4). Only at the end of the first year of life, the appearance of highly oxygen-sensitive butyrate producers – including *Faecalibacterium*, the hallmark of the healthy adult gut ecosystem[17]– was observed (data not shown). Microbial production of butyrate is of key importance to create and maintain the anaerobic conditions that characterize a healthy, adult colon environment[18].

### The effect of external factors on infant gut microbiota maturation

Although maturation of the infant gut microbiota was identified to be a largely unidirectional process, occasional transient regression towards a preceding gut microbiota maturation stage could be observed (Figure 3a). Hypothesizing maturation stage regression to be associated with disease or medical interventions, we developed an ecosystem maturation index per sample based on presence/absence of genera belonging to the BaBel average top 15. As discussed above, we ranked each genus according to its order of appearance along the timeline of an infant’s ecosystem maturation process. Next, genera were attributed an overall cohort rank (1 to 10, Figure 2b) based on their median order of appearance across individual infants. A samples’ maturation index was calculated by averaging the ranks of the present genera (relative abundance >0.5%, Figure 3b). We identified three time points (events) displaying a lower maturation score than expected (*i.e.* outside the 95% CI of the regression of the maturation score) concurring with a regression in maturation stage (Figure 3a). A first event (E1; infant S004 at day 163, regression from maturation stage B to A) coincided with the end of a seven-day oral antibiotic treatment (day 155 to 161; amoxicillin with the adjuvant clavulanic acid, a β-lactamase inhibitor) for a urinary tract infection. After treatment initiation, *Streptococcus* became the predominant genus, falling back below detectable levels two days after the last dose of antibiotics (Figure 3c). Multivariate analysis on the extended BaBel dataset (including all eight infants) identified *Streptococcus* as the genus most significantly increased in abundance during antibiotic treatment (n=299, dbRDA using all covariates, adjusted R^2^=0.12, FDR<0.05; n=303, MaAsLin2 testing all covariates on all genera, FDR=0.0011; Supplementary Table 1h; Figure 2a). Genera with lowered proportional abundances upon amoxicillin treatment included *Bifidobacterium* and *Veillonella*, both decreasing below detection limits and reappearing after less than 18 and 6 days after cessation of treatment, respectively (Figure 3c). After the disappearance of *Streptococcus*, *Escherichia* was the first genus to re-establish, becoming the most dominant member of the gut microbiota less than 2 days after the last dose of amoxicillin (Figure 3c). These observations confirm the status of oxygen-tolerant genera as pioneering colonizers in primary succession as well as secondary colonization following antibiotic treatment-associated ecosystem disruption, with gut microbiota maturation stage regression probably associated with an imbalance in colon oxygen homeostasis[19] (Figure 3a,c). Of note, two other infants (S003, days 353-359; and S010, days 214-220) also received amoxicillin (without clavulanic acid), in both cases prescribed to treat an ear infection. However, only less pronounced microbiome alterations were observed upon treatment, possibly due to the absence of an adjuvant or to the fact that the infants’ microbiota had matured to the potentially more stable C maturation stage. The second event (E2; infant S009 at day 251, regression C to B) coincided with an untreated *Cryptosporidium* infection (days 248- 250), accompanied by fever and diarrhoea, which was characterized by a observed rise in relative abundances of *Bifidobacterium* and *Streptococcus,* while the other genera decrease (Figure 3 a,b,d). E3 (S011, days 13-21) co-occurred with the start of a period of severe constipation in infant S011 (Figure 3e). While the baby’s first samples taken at days 6 and 7 were classified within the infant-specific maturation stage D (Figure 3a), a transition to the *Bifidobacterium*-dominated B maturation stage could be noted on days 13, 17, and 21. During the period following maturation stage regression, infant S011 suffered from recurrent episodes of severe constipation, including three periods of 6 to 9 days without bowel movement (defecation on days 32, 40, 41, 47, and 53). However, from day 32, the infant’s faecal microbiome returned to the maturation state D classification.

**Figure 3.**
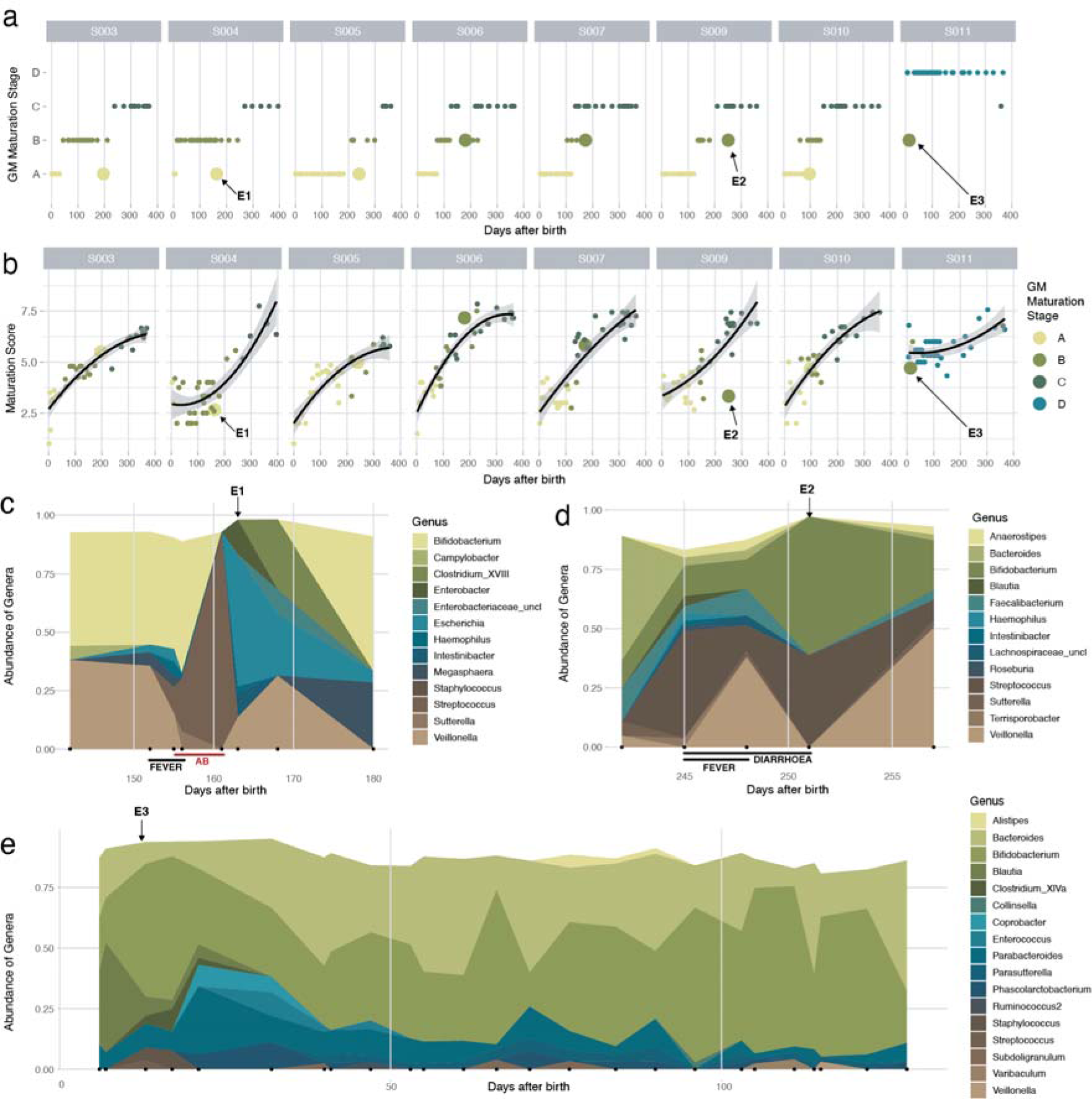
The effect of external factors on the infant gut microbiome. (a) Succession of the gut microbiota maturation stages over time, including all 303 time points from the BaBel dataset. Time points representing a return to a previous gut microbiota maturation stage (after at least 2 samples in the next gut microbiota maturation stage), are represented with larger dots. (b) The change in maturation score of the samples over time. The maturation score was calculated by averaging the ranks (based on their order of appearance) of the present genera in every sample. The black line represents the quadratic regression with 95% confidence interval (all p-values of the quadratic fits < 0.0002). Three events, for which the succession goes back to a previous gut microbiota maturation stage (shown in 4a) and the maturation score drops (outside the confidence interval) are indicated with the arrows. (c) Changes in bacterial abundance during the antibiotic event in infant S004 (“E1” at day 163, abundances >0.02 shown). The red line indicates the duration of the treatment (7 days) with antibiotics (amoxicillin and clavulanic acid). (d) Changes in abundance during a *Cryptosporidium* infection in infant S009 (“E2” at day 251, abundances >0.02 shown). (e) Changes in abundances in the first half year in infant S011 (“E3”, at days 13-21, abundances >0.05 shown).

### Transition of the infant gut microbiota maturation towards an adult configuration

To evaluate gut microbiota maturation during the first year of life in terms of ecosystem transition towards an adult configuration, we mapped the microbiome composition of the infant samples onto the background of inter-individual variation as observed in the Flemish Gut Flora Project (FGFP) population cohort (n=1,106; Figure 4). Previously, using DMM-based community typing[7], genus-level compositional differentiation of the adult microbiome in the FGFP has been shown to revolve around four enterotypes[20] – prevalent, non-discrete microbiome constellations that can be identified reproducibly across datasets[20–22]. Having aligned not only DNA extraction and sequencing methods, but also analytical procedures with the FGFP protocols[23], we observed the faecal microbiomes of Flemish infants to differ substantially from those obtained from adults inhabiting the same region (permutational MANOVA Adonis test, n=1,407, R^2^=0.30, p-value=0.001; Figure 4b,c,d). All infant samples were however classified as *Bacteroides*2 (Bact2) communities (Supplementary Table 1i; Figure 4a,b) – a recently described low-diversity/low cell density constellation characterised by high *Bacteroides* and low *Faecalibacterium* proportional abundances. Bact2 communities have previously been linked to loose stools[21], inflammation[21] and reduced wellbeing[24], and have been hypothesized to reflect an ecosystem dysbiosis[20, 21, 25]. The similarities of infant microbiota constellations to adult dysbiotic states, as previously noted[6], are likely attributable to convergences between primary (ecosystem development) and secondary (perturbation recovery) succession[6, 26]. Like in adult dysbiosis, the infant gut ecosystem has been reported to display low colonization resistance[15, 27], exemplified by the frequency of gastrointestinal infections reported in the present cohort - with Babel infants experiencing on average two (range = [0:3]) episodes of diarrhoea during the first year of their life - and beyond[28]. At the same time, a shift in the infant microbiota composition towards a more adult-like configuration could be observed over time. When comparing the microbiota composition of BaBel age bins [0:3, 3:6, 6:9, and 9:12 months] with the FGFP population cohort, effect sizes in microbiome variation were observed to decrease with increasing infant age (permutational pair-wise MANOVA Adonis test, n=[1206:1204:1153:1159], R^2^=[0.228:0.221:0.085:0.067]; FDR<0.01; Supplementary Table 1j; Figure 4c). Moreover, a detailed analysis of DMM clustering result identified six samples from three infants taken in the last month of their first year having a non-zero probability of not belonging to the Bact2 community type (probability range=[4.34e-6:1.20e-14]; Supplementary Table 1i). In all samples, the observed transition towards a more adult microbiome constellation was accompanied by an increase in observed genus richness over time– although adult richness was not reached (infant age bins vs adults, KW and phD tests, n=[1207:1205:1154:1160], r=[- 0.52:-0.47:-0.32:-0.32], FDR<0.05, Supplementary Table 1k; Figure 4e).

**Figure 4.**
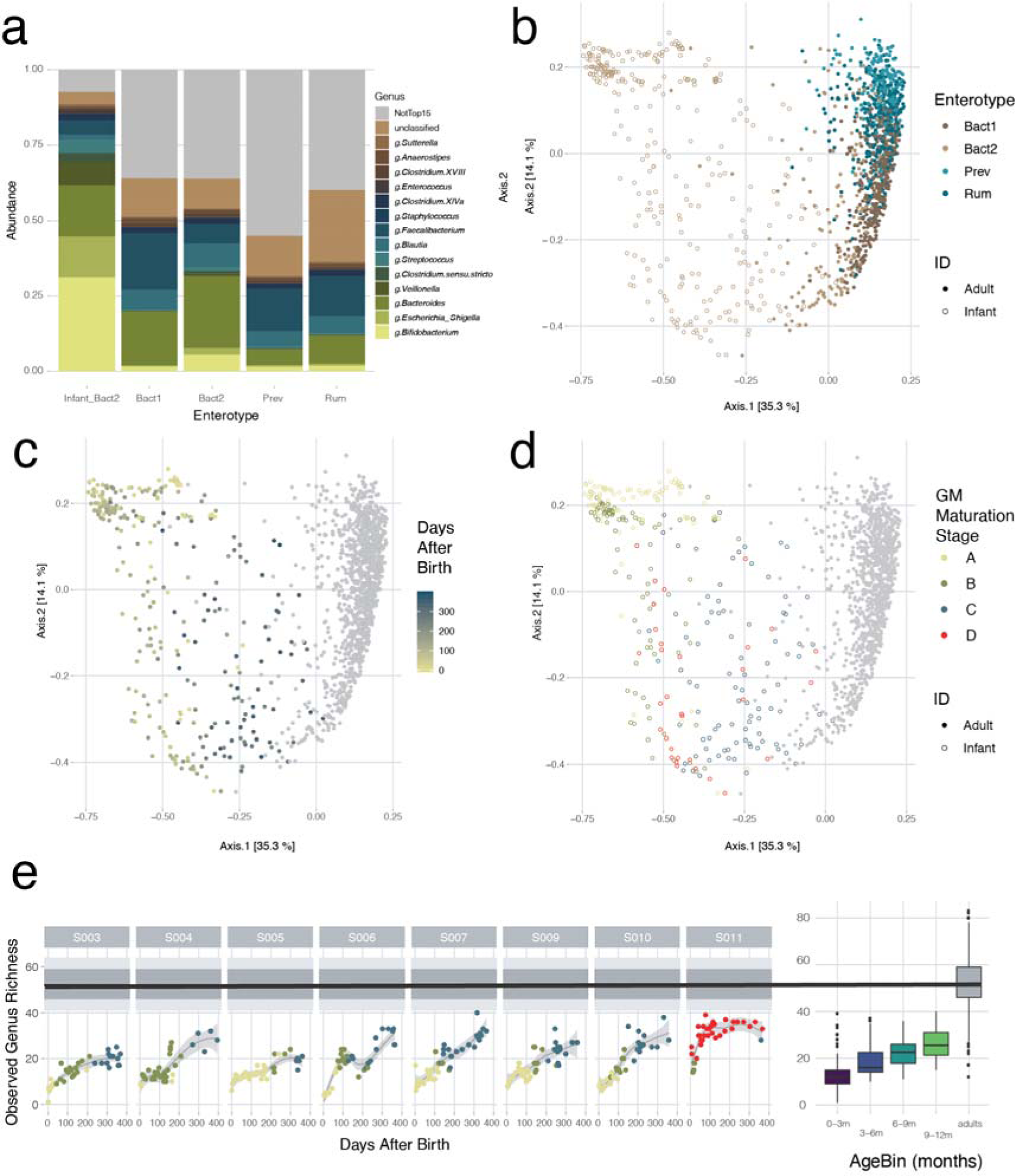
Projection of the infant sample to adult samples of the Flemish Gut Flora Project (FGFP) dataset. (a) Barplots showing the average relative abundances of the top 15 most common bacterial genera of the infant samples and the adult samples, per enterotype. (b) Projection of the infant samples to the adult FGFP dataset, visualized on a principle coordinate analysis (PCoA, Bray-Curtis dissimilarity), colored for enterotype, (c) colored for time after birth (for the infant samples), (d) colored per gut microbiota maturation stage. (e) Observed genus-level richness over time of the BaBel dataset (Loess smoothing), compared to the observed genus level richness of the FGFP dataset (black line is the median, dark gray area represents the 25-75 IQR and the light gray area represents the 10-90 IQR). On the right side, the boxplots represent the genus level richness for the different infant age bins, compared to the adult FGFP dataset. The body of the box plots represent the first and third quartiles of the distribution and the median line.

## CONCLUSION

We show that maturation of gut microbiota can be captured in a series of transitions that remain conserved across the BaBel infants – both on the community/gut microbiota maturation stage level as in order of appearance of prevalent genera. Throughout the first year of life, successional colonization of the gut microbiota results in a shift from a low richness, oxygen tolerant community dominated by pioneering colonizers such as *Escherichia* to a more diverse community comprising anaerobic butyrogens such as *Faecalibacterium* – with butyrate being a key metabolite in maintenance of colonic hypoxia[18]. Our analyses confirm previously reported similarities between the infant microbiota and adult dysbiosis[6, 29, 30] likely due to shared features of primary and secondary succession. While temporary regression following ecosystem-disrupting events such as infection or antibiotic treatment can be observed, the microbiota of all studied infants matured to a more adult-like constellation over the first year of their life, as reported before[31]. Given the similarities observed between primary succession and secondary colonization upon disruption, careful dissection of the succession events characterizing gut ecosystem maturation could pave the way for the development of mimicking biotherapeutic strategies in adult microbiome modulation.

## METHODS

### Sample collection

Between 2013 and 2017, stool samples of eight Belgian healthy infants, *i.e.* the BaBel infants, were collected starting from birth at a frequency of 2-3 samples per week (Supplementary Table 1a). Samples were kept at -20°C freezers at the participants’ homes and every three months transported to our laboratory on dry ice, where they were stored at -80°C until further analysis. Every time a sample was collected, the parents completed a questionnaire containing information about the date, consistency of the stool (aqueous/soft/solid), diet (breastmilk/formula milk/vegetables/fruit), clinical signs or disease (diarrhoea/vomiting/fever/…), and the location of the infant when the sample was taken (at home/day care/holiday location/…). All infants were vaginally born, the mothers did not take antibiotics during pregnancy or delivery, and no complications during pregnancy were reported. The histo-blood group antigen (HBGA) specificities (ABO group antigens, Lewis antigens, FUT2 and FUT3 genotype) were determined as described before[32], from a saliva sample from each infant collected at the end of the study period. For the investigation of the overall effect of metadata on the microbiome composition, only covariates present in at least three infants were used (infant ID, time after birth, presence of furry pets, secretor-status, Lewis antigens, ABO blood group, diet pattern (BreastOnly/NoSolid/Solid), consistency, diarrhoea, fever, respiratory illness and other general sickness signs, painkillers, antibiotics and day care).

### Sample selection

To study the longitudinal dynamics in the gut microbiome, 21 stool samples from predefined days 0, 3 7, 10, 15, 21, 30, 45, 60, 75, 80, 105,120, 150, 180, 210,240, 270, 300, 330 and 360 were selected from each of the eight infants (Supplementary Figure 1). When an infant showed clinical signs at any of these time points, we selected the closest available sample without clinical signs present, or this time point was excluded. In total, we included 159 samples at predefined timepoints, of which 17 felt together with clinical signs (and were not replaceable by a close timepoint with no signs) and 142 did not fall together with clinical signs (Supplementary Table 1a, Supplementary Figure 1). In addition, we selected 144 additional samples *ad hoc* from before, during and after specific external events to study how they influence the gut microbiome (events included vaccination history, type of food consumed, occurrence of diseases, use of antibiotics, use of pre- or probiotics; Supplementary Figure 1).

### 16S rRNA gene library preparation and sequencing

Bacterial profiling was carried out as described by Falony and colleagues[23]. Briefly, nucleic acids were extracted from frozen faecal aliquots using the RNeasy PowerMicrobiome kit (Qiagen). The manufacturer’s protocol was modified by the addition of a heating step at 90°C for 10min after vortexing and by the exclusion of DNA-removal steps. Microbiome characterization was performed as previously described[33], in short, the extracted DNA was further amplified in triplicate using 16S primers 515F(5’-GTGYCAGCMGCCGCGGTAA-3’) and 806R(5’-GGACTACNVGGGTWTCTAAT-3’) targeting the V4 region, modified to contain a barcode sequence between each primer and the Illumina adaptor sequences to produce dual-barcoded libraries. Deep sequencing was performed on a MiSeq platform (2x250PE reads, Illumina). All samples were randomized and negative controls were taken along and sequenced.

### Sequenced read analysis

After demultiplexing with sdm as part of the LotuS pipeline[34] without allowing for mismatches, fastq sequences were further analysed per sample using DADA2 pipeline (v. 1.6)[35]. Briefly, we removed the primer sequences and the first 10 nucleotides after the primer. After merging paired sequences and removing chimeras, taxonomy was assigned using formatted RDP training set ‘rdp_train_set_16’. The decontam[36] R package was used to remove contaminating Amplicon Sequencing Variants (ASVs) using the frequency prevalence method(Supplementary Table 1l). After quality control steps, the ASV table contained on average 46,330 reads per sample (range = 15427-131451). In total 197 ASVs were obtained all belonging to the kingdom Bacteria. No Archaea were detected. All samples were rarefied to 14,668 reads per sample and ASVs with an overall relative abundance <0.0001 were removed. From three samples (S009-1, S004-1 and S010-1), of three different infants the first sample taken, we were not able to extract enough DNA to be amplifiable.

### Statistical analyses

All statistical analyses were performed and visualized in R (http://www.R-project.org) using the ggplot2[37], phyloseq[38], synchrony[39], DirichletMultinomial[40], dunn.test[41] and vegan[42] packages. To test median differences between two or more groups of continuous variables, Mann-Whitney U test and Kruskal-Wallis (KW) test were performed respectively. The KW test was always followed by post hoc Dunn’s (phD) test for all pairs of comparisons between groups. Multiple testing correction was performed where appropriate using the Benjamini-Hochberg procedure (FDR-adjustment set at <0.05).

### DMM clustering to identify the colonization stages

To determine the stages of the colonization process, a Dirichlet Multinomial Mixtures (DMM) based approach was followed, as described by Holmes *et al.*[7] using the DirichletMultinomial[40] R package on the genus level (rarefied) read matrix (n=303). The optimal number of stages was determined based on Bayesian information criterion (BIC) and the mean probability for the samples to belong to the assigned Dirichlet component was on average 0.99 (median=1, stdev=0.05, Supplementary Table 1b).

### Determination of the order of appearance of the top genera

Per infant, the 15 most abundant genera (present in more than 3 infants) were ranked based on the first timepoint in which they were present (with an abundance >0.5%). Rankings were scored using Kendal w-test using the R function *kendall.w* of the synchrony[39] package with 10,000 permutations. A final order of appearance was set, based on the order of the medians of the ranks per infant. Finally, a maturation score was calculated for every sample by averaging the ranks of the genera weighted by the presence or absence of that specific genus. Growth rates (GR) of the different genera were calculated from the predicted generation times (GT=1/GR), as published before[43].

### Alpha and Beta diversity

Alpha-diversity (richness and Shannon diversity) and beta-diversity indices (Bray-Curtis dissimilarity) were calculated by using the phyloseq[38] package. Ordinations were visualized on a principle coordinate analysis (PCoA) using Bray-Curtis dissimilarity. The univariate effect of the metadata variables on the first two axis of the ordination are determined using *envfit* function of the vegan[42] package (univariate distance-based redundancy analysis (dbRDA)) and plotted as arrows on the PCoA (InfantID was excluded for clarity). Community-level differences between groups were tested with Adonis non-parametric test of the vegan[42] package. If more than two groups are compared, a post-hoc Adonis test was used in a pairwise way, correcting for multiple testing.

### Multivariate analysis of the effect of metadata variables on microbial composition

To investigate which metadata covariates contribute to the variation in microbiota community, dbRDA was performed on genus level (Bray Curtis distance), using the *capscale* function in the vegan[42] R-package. Covariates found to significantly contribute to the ordination outcome were further implemented in forward model selection on dbRDA using the *ordiR2step* function in the vegan[42] package, to determine the non-redundant cumulative contribution of metadata variables to the variation (stepwise dbRDA). To test the effect of metadata variables on specific genera, the same approach as previously described was followed by first pruning the community to only contain the genus of interest (for each of the top 15 genera), followed by dbRDA on the Euclidean distances measured on the abundances of that genus and forward model selection as described above, constraining for infant ID. To confirm results from the previous step, MaAsLin2[44] was used, which performs boosted additive general linear models to discover associated between metadata and the relative taxonomic abundances (default settings). Note, that only for the dbRDA four samples were excluded for which consistency was unknown (n=299).

### Projection to the adult FGFP dataset

Enterotypes of the infant samples were computed against a background of adult non-disease-associated microbiomes (FGFP dataset, genus-level abundance matrix, n=1,106) by DMM clustering using the DirichletMultinomial package as described by Holmes *et al*.[7] Samples were rarefied to 10,000 reads. To avoid interference by non-independent samples, enterotyping was performed iteratively on one randomly-selected sample of each infant against the FGFP background (n=42 enterotyping rounds). The optimal number of Dirichlet components based on BIC was four in all iterations, and the clusters were named *Prevotella*, *Bacteroides*1, *Bacteroides*2, and *Ruminococcaceae* as described before[20].

## DECLARATIONS

### Ethics approval

The study was approved by the IRB at KU Leuven (ML8699, S54745, B322201215465).

### Consent for publication

Not applicable.

### Availability of data and materials

16S sequencing data used in this study is available at the European Nucleotide Archive (ENA, https://www.ebi.ac.uk/ena, PRJEB40751, <u>not accessible for public yet</u>). The code to perform analysis and make figures starting from the ASV abundance table <u>will be</u> made available at https://github.com/Matthijnssenslab/BabyGut16S/.

### Authors’ contributions

The study was conceived by JM, JR and MVR. Experiments were designed by JM, JR, LB, MZ and RT. Sampling was set up and carried out by CS, JM, KCY, KF, LB and WD. Experiments were performed by DJ, LB, LVE and LR. LB, MIP and RT performed the bio- informatics analyses of the sequenced reads. Statistical analyses were designed and performed by GF, LB, MV-C, SV-S and WD. LB, JM, GF, SV-S, MIP, MZ and WD drafted the manuscript. All authors revised the article and approved the final version for publication.

## Acknowledgements

We would like to thank all participating infants and parents for their contribution. We thank Dr. Johan Nordgren for the characterisation of the saliva samples.

## Authors’ information

Correspondence should be addressed to JM and JR.

## SUPPLEMENTARY FIGURES WITH LEGENDS

**Supplementary Figure 1.**
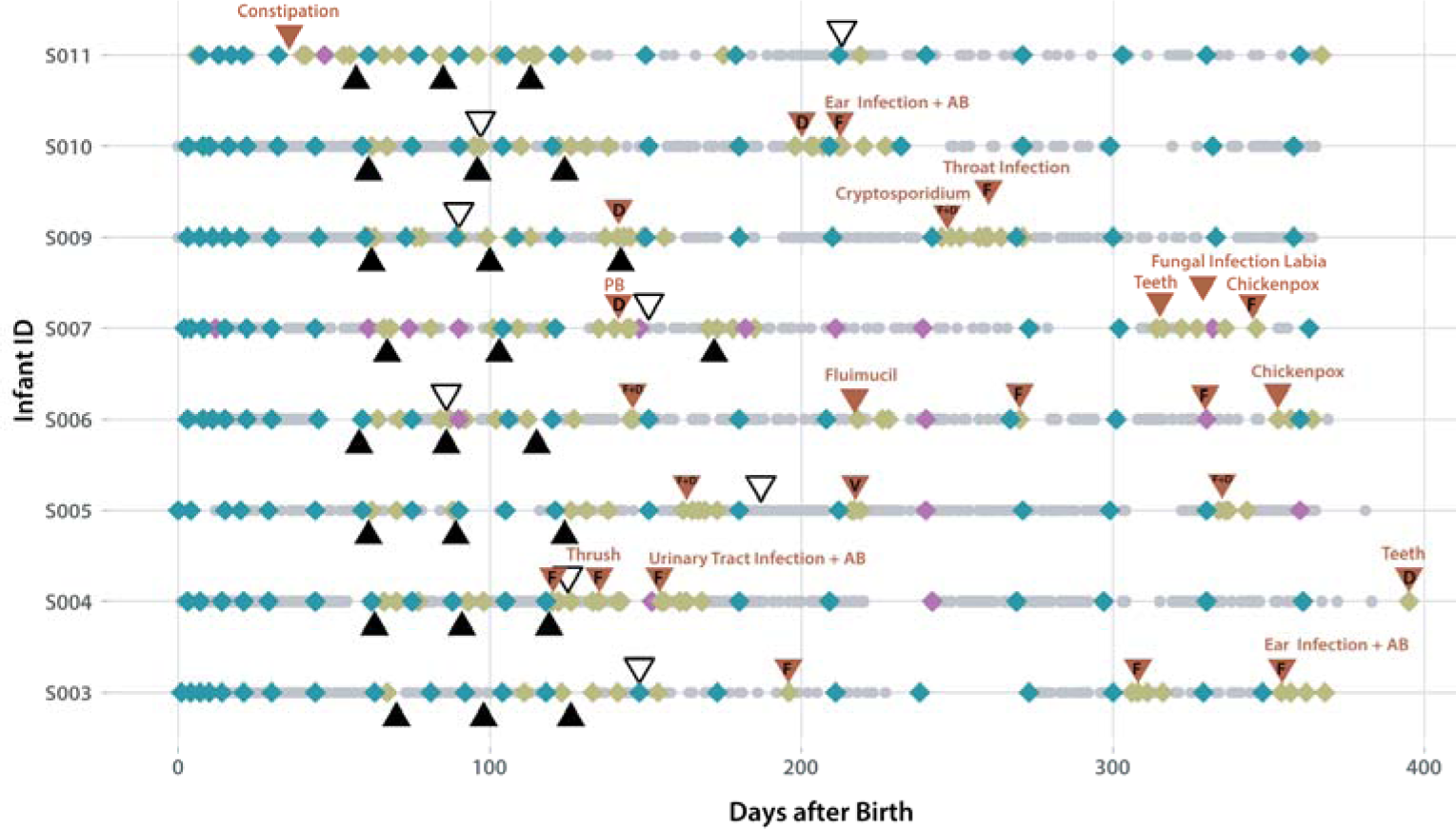
Overview of the collected and selected samples per infant. Grey dots 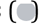: All samples collected by the parents of the enrolled infants Blue diamonds 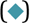: Samples selected for the study of the longitudinal dynamics at predefined timepoints with no clinical signs (n = 142) Purple diamonds 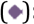: Samples selected for the study of the longitudinal dynamics at predefined timepoints with clinical signs (n = 17) (See supplementary Table 1a for the signs) Green diamonds 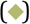: Additional *ad hoc* selected samples at specific external events (n = 144) Black filled triangles 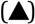: Three vaccinations events in every infant Black open triangles 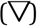: Day care entrance Red triangles 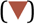: External events around which extra samples were selected. Abbreviations: fever (F), diarrhoea (D), vomit (V), antibiotics (AB), Probiotics (PB)

**Supplementary Figure 2.**
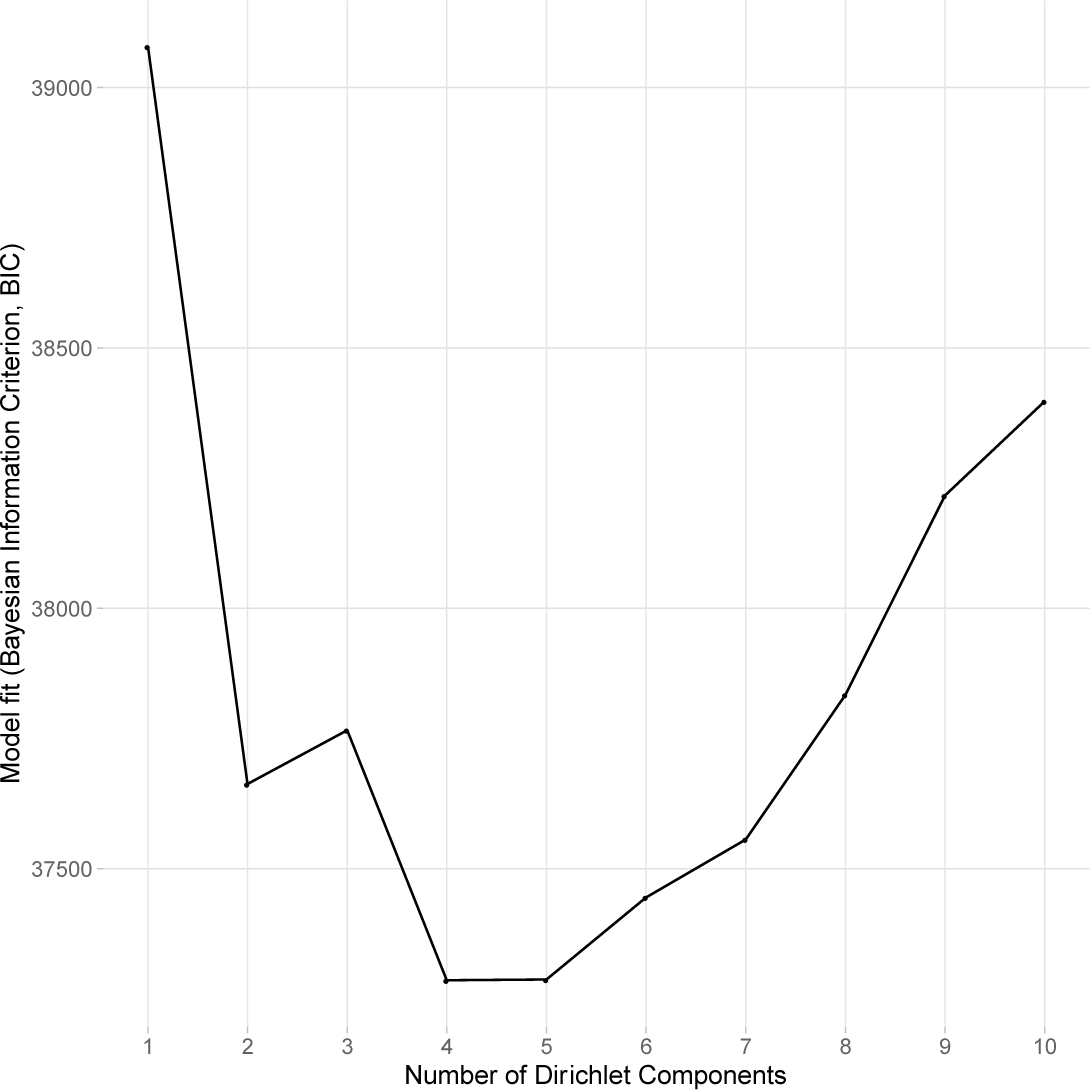
Determination of the optimal number of clusters in the DMM approach. Identification of the optimal number of Dirichlet components in the BaBel dataset (N=303) based on the Bayesian Information Criterion (BIC). The optimal number of clusters is four (minimum BIC= 37285.6).

**Supplementary Figure 3.**
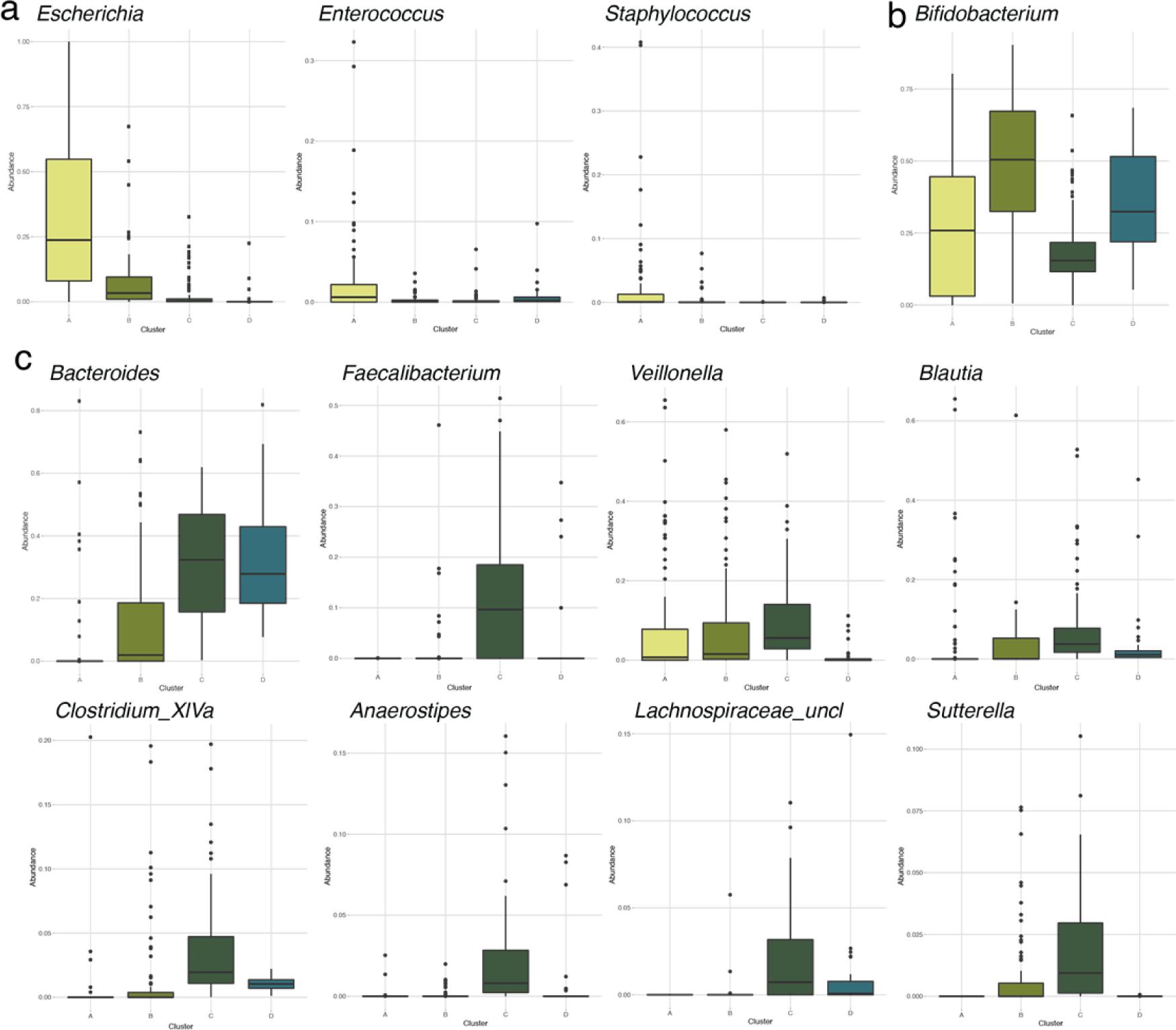
Most abundant genera that are differentially abundant per gut microbiota maturation stage determined using DMM clustering. (a) Distribution of the relative abundances of the most abundant genera in gut microbiota maturation stage A, that are significantly more abundant in maturation stage A than in B and C. (b) Distribution of the relative abundances of the most abundant genera in gut microbiota maturation stage B, that are significantly more abundant in maturation stage B than in A and C. (c) Distribution of the relative abundances of the most abundant genera in gut microbiota maturation stage C, that are significantly more abundant in maturation stage C than in A and B. (n = 303, KW with phD test, r > 0.3, FDR < 0.05; Supplementary Table 1d)

**Supplementary Figure 4.**
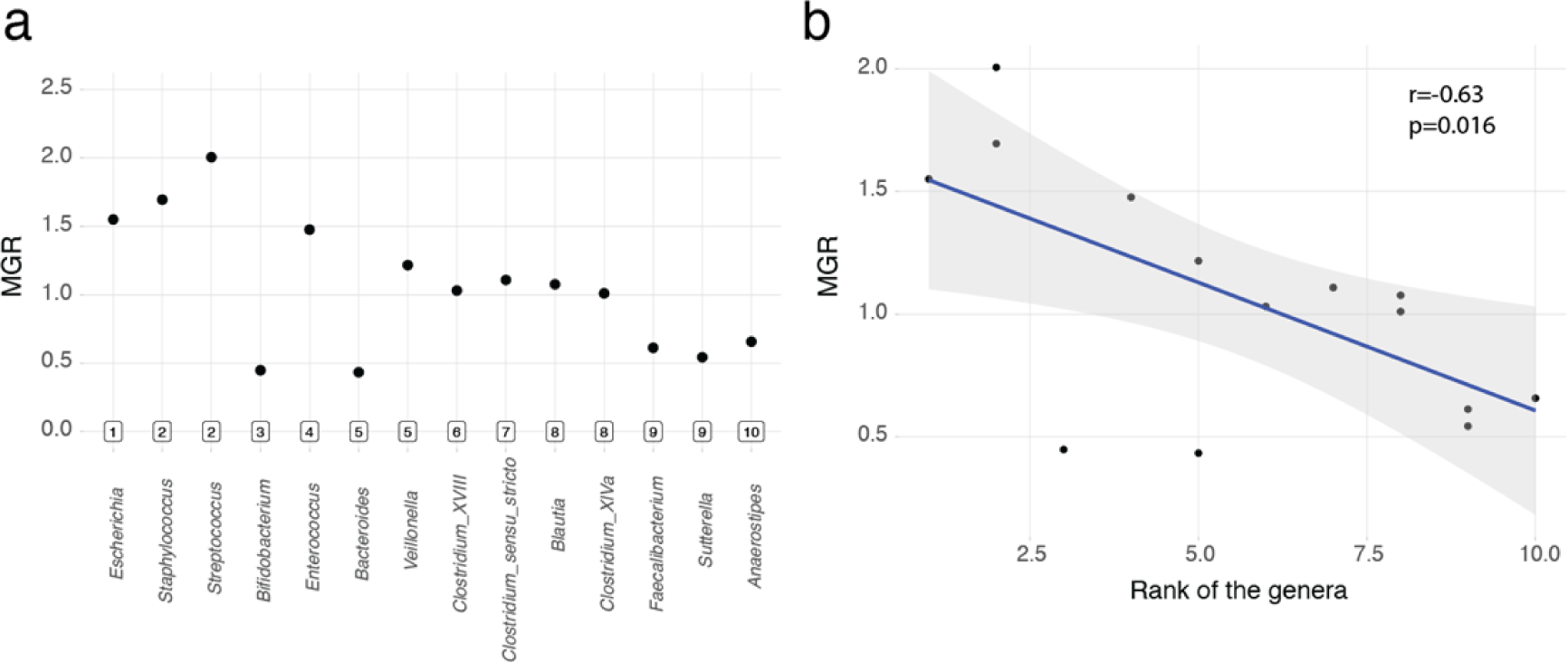
Average predicted growth rates for the top genera of the infant gut. (a) The maximum growth rates (MGR) of the top 15 most abundant genera in the infant gut, ordered by their rank of appearance, calculated like reported before[43]. Note that for one genus, *Lachnospiraceae* unclassified, no growth rate could be obtained. (b) Negative correlation between the ranks of the top genera and their growth rates (Pearson correlation coefficient, n = 14).

